# Spontaneous behavioural recovery following stroke relates to the integrity of sensory and association cortices

**DOI:** 10.1101/2022.02.18.481083

**Authors:** Joseph Y. Nashed, Kaden T. Shearer, Justin Z. Wang, Yining Chen, Elise E. Cook, Allen A. Champagne, Nicole S. Coverdale, Juan Fernandez-Ruiz, Shirley I. Striver, Jason P. Gallivan, Douglas J. Cook

## Abstract

Stroke is a devastating disease that results in neurological deficits and represents a leading cause of death and disability worldwide. Following a stroke, there is a degree of spontaneous recovery of function, the neural basis of which is of great interest among clinicians in their efforts to reduce disability following stroke and enhance rehabilitation. Conventionally, work on spontaneous recovery has tended to focus on the neural reorganization of motor cortical regions, with comparably little attention being paid to changes in non-motor regions and how these relate to recovery. Here we show, using structural neuroimaging in a macaque stroke model and by exploiting individual differences in spontaneous behavioural recovery, that the preservation of regions in the sensory and association cortices predict animal recovery. To characterize recovery, we performed a clustering analysis using Non-Human Primate Stroke Scale (NHPSS) scores and identified a good versus poor recovery group. By comparing the preservation of brain volumes in the two groups, we found that brain areas in integrity of brain areas in parietal, temporal and somatosensory cortex were associated with better recovery. In addition, a decoding approach performed across all subjects revealed that the preservation of specific brain regions in the parietal, somatosensory and medial frontal cortex predicted recovery. Together, these findings highlight the importance of the sensory and association regions in spontaneous behavioural recovery, supporting the notion that the ‘sensory retraining’ of impairments following stroke is critical to recovery and motor function.

## Introduction

Stroke is a major public health issue and represents a leading cause of adult disability (Virani et al., 2020; Zorowitz, Chen, Tong, & Laouri, 2009). Though stroke is known to cause deficits in several neurological domains, the greatest hurdle hindering functional independence in those affected by stroke is the resulting impairment in motor function (Bernspang, Asplund, Eriksson, & Fugl-Meyer, 1987; Lai, Studenski, Duncan, & Perera, 2002). Following stroke, there is a degree of spontaneous recovery that results from the intrinsic rewiring of salvaged brain regions (Cramer, 2008; Feydy et al., 2002; Ward, Brown, Thompson, & Frackowiak, 2003; Wei et al., 2013). The majority of this spontaneous recovery generally occurs within 1 month post-stroke, but with significant improvements continuing for up to 3 months to a year (Dobkin, 2003; Wade, Langton-Hewer, Wood, Skilbeck, & Ismail, 1983). Characterizing this recovery process with precision has proven challenging, as there exists considerable variability in both the degree and timeline of recovery across stroke survivors. While some of this variability relates to the tools used to assess motor behaviour, (Coderre et al., 2010; Dukelow et al., 2010; Gladstone, Black, Hakim, Heart, & Stroke Foundation of Ontario Centre of Excellence in Stroke, 2002; Wityk, Pessin, Kaplan, & Caplan, 1994) a greater source of this variability likely arises due to the fact that recovery is dependent on the brain region affected by the stroke.

Given that an improvement in patients’ motor control is the defining hallmark of spontaneous behavioural recovery, it is not surprising that studies have tended to focus on the role of motor-related brain areas (e.g., primary and premotor cortex) in driving this recovery (Bolognini, Russo, & Edwards, 2016; Edwards, King, Buetefisch, & Borich, 2019). However, in addition to the ability to generate descending motor commands, recent theories of motor control strongly emphasize the importance of sensory feedback in guiding successful motor actions. In optimal feedback control models (Scott, 2004; Todorov & Jordan, 2002), for instance, the generation (and correction) of movement involves the real-time manipulation of sensory feedback to achieve a desired motor goal (Kurtzer, Pruszynski, & Scott, 2008; Nashed, Crevecoeur, & Scott, 2012, 2014; Nashed, Kurtzer, & Scott, 2015; Pruszynski et al., 2011; Pruszynski, Kurtzer, & Scott, 2008; Yang, Michaels, Pruszynski, & Scott, 2011). Consistent with this framework, animal studies have illustrated that focal lesions to somatosensory areas result in impaired motor behaviour, as well as deficits in motor learning and the generation of fine motor skills (Hiraba, Yamaguchi, Satoh, Ishibashi, & Iwamura, 2000; Lin, Murray, & Sessle, 1993; Pavlides, Miyashita, & Asanuma, 1993; Xerri, Merzenich, Peterson, & Jenkins, 1998; Yao, Yamamura, Narita, Murray, & Sessle, 2002). However, despite the critical importance of sensory feedback to motor control, human and non-human primate research has tended to focus largely on motor recovery following stroke without accounting for the important role of sensory processing (Edwards et al., 2019).

Here we investigated the neural substrates mediating spontaneous behavioural recovery by studying natural variation in the recovery of thirty-one cynomolgus macaques following stroke. This previously validated animal model of stroke was selected due to the phylogenetic similarities in anatomy, physiology and behaviour between cynomolgus macaques and humans (Cook & Tymianski, 2012). Each macaque underwent a transient 90-minute Middle Cerebral Artery Occlusion (MCAO) and had their motor functions tested daily, using the Non-Human Primate Stroke Scale (NHPSS), for 30 days following the occlusion (Roitberg et al., 2003). At 30 days post-stroke, we then performed whole-brain structural neuroimaging using T2-weighted MRI images. We found, through a data-driven clustering of animals into groups that exhibited good versus poor behavioural recovery, that the remaining cortical volumes of sensory-and association-related brain areas—and not primary and secondary motor areas—was associated with good recovery. Notably, this same association was not observed when instead using stroke volume as a predictor. Together, our findings suggest that it is the integrity of sensory and association cortices that determines the degree of spontaneous behavioural recovery following stroke.

## Material and methods

### Animal preparation

Data was obtained from 31 cynomolgus macaques (Macaca fascicularis) with an average weight of 4.13 ± 1.1 kg (range 3.6 to 5.3 kg). All the surgical and experimental procedures were carried out in accordance with the Canadian Council of Animal Care policy on the use of laboratory animals and approved by the Animal Use Subcommittee of Queen’s University Council on Animal Care.

### Stroke model

Animals were anesthetized (Isoflurane 1.0–1.6%), intubated and ventilated. Non-invasive monitoring included blood pressure by leg cuff, end-tidal CO2, O2 saturation, ECG and temperature by rectal probe. Temperature was maintained (37 ± 0.5 °C) by a heating blanket. A femoral arterial line was used to monitor blood pressure and blood gases. A middle cerebral artery occlusion (MCAO) was performed using a right pterional craniotomy (Fig. 1a). The sylvian fissure was carefully dissected to expose the right middle cerebral artery (MCA), which was subsequently occluded with a 5mm titanium micro aneurysm clip along the M1 subdivision of the MCA just proximal to the orbitofrontal branch (Fig. 1b). Animals were then transferred to the scanner for Magnetic Resonance Imaging (MRI) to confirm occlusion by magnetic resonance angiography (MRA). Ischemia duration was 90.0 ± 1.1 min across the 31 animals. After occlusion, the incision was reopened, and the aneurysm clip was removed in order to restore blood flow. The craniotomy was then irrigated with 0.9% NaCl solution and the dura, temporalis muscle, fascia and skin were closed. The resulting MCAO resulted in a well-defined stroke in all animals.

**Figure 1:**
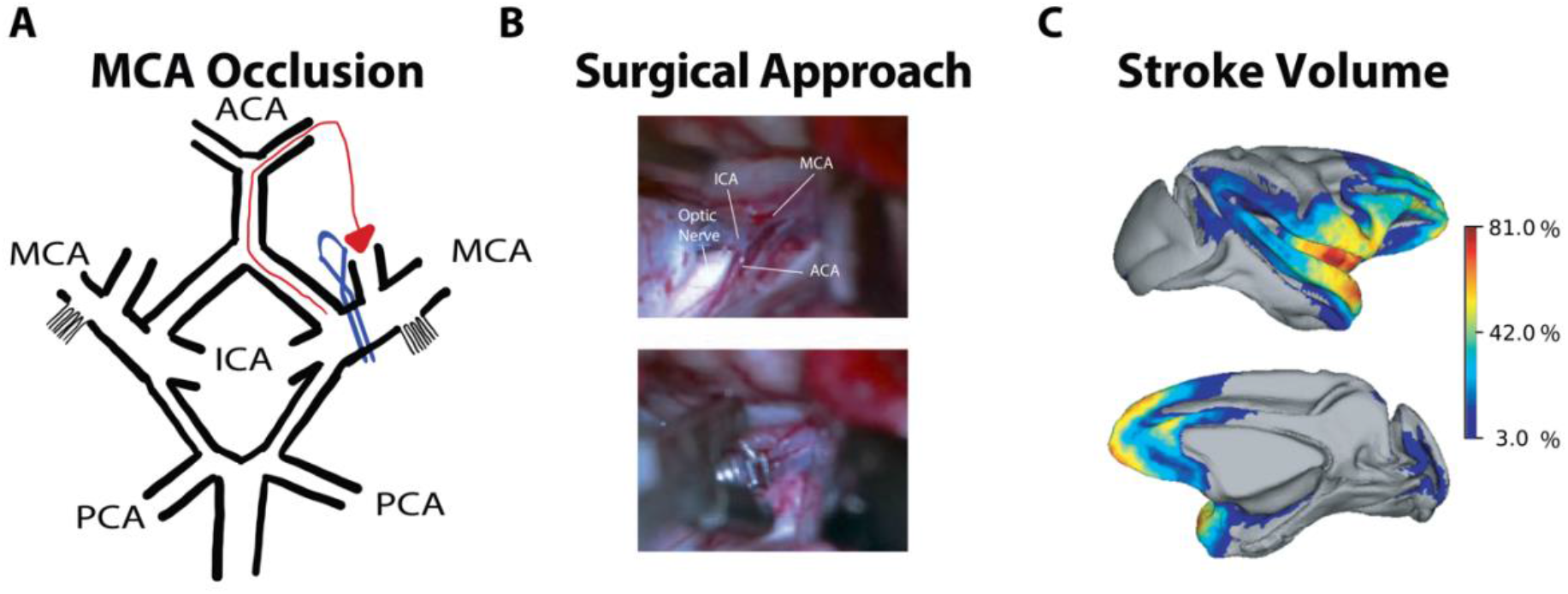
Cynomolgus macaque model of right middle cerebral artery occlusion (MCAO). **A)** Schematic of surgical method of MCAO illustrating placement of aneurysm clip (blue) and collateral blood flow through the anterior and leticulostriate cerebral collaterals. **B)** Surgical methodology, exposing the M1 segment of the Middle Cerebral Artery (MCA) (Top) and placement of a #5 Sundt AVM clip to the M1 segment of the MCA (Bottom). **C)** Resultant stroke area across all animals. Mean percent affected area following MCA occlusion in 31 monkeys. The legend denotes the percent of all animals with areas affected. ACA = Anterior Cerebral Artery; ICA = Internal Carotid Artery; PCA = Posterior Cerebral Artery

### Neurological assessment

Neurological function was assessed prior to and following MCAO using the Non-Human Primate Stroke Scale (NHPSS) at 1,2,3,4,5,7,14,21,28 and 30 days. The NHPSS is akin to the NIH Stroke Scale in humans and is used to quantify stroke severity based on measures aspects of consciousness, grasp reflex, extremity movement (upper limbs and lower limbs), neglect, hemianopia, and facial weakness (Cook, Teves, & Tymianski, 2012; Roitberg et al., 2003). The NHPSS is composed of 11 domains and it yields a total score of 41 points, where 0 corresponds to normal behaviour, and where 41 corresponds to severe bilateral neurological impairment. Previous experiments in cynomolgus macaques with 90 min MCAO showed an initial increase in NHPSS scores, similar to human patients, and gradually declined and plateaued between 14 and 30 days, indicating that the behavioural impairment had become stable at one-month post stroke (Cook et al., 2012; Nacu et al., 2016).

### MRI Image Acquisition

Animals underwent three scanning sessions, 1) pre-stroke (baseline), 2) <1h post-stroke (stroke confirmation), and 3) 30 days post-stroke. All data were acquired on an actively shielded 3 Tesla Siemens Trio scanner with a 32-channel head coil at the Queen’s University Centre for Neuroscience Studies. For the acquisition of MRI images, animals were intubated, and anesthetized (Isoflurane 1.0–1.6 %, O_2_ flow rate of 2 L/min) throughout scanning. The induction of anesthesia was performed in the same way as in the surgical procedure with a mixture of ketamine (7.5 mg/kg) and dexmetatomadine (0.05 mg/kg). Baseline MRI acquisitions were made 14 days prior to MCAO. Confirmation of vessel occlusion post stroke was made using a high-resolution MRA sequence; TR = 39 ms; TE = 7.33 ms, flip angle = 15°, matrix = 448 × 448, FOV 112 × 112 mm^2^ and final voxel size = 0.3 × 0.3 × 1 mm^3^. Thirty days post stroke scans consisted of a high-resolution T2-weighted Fast Spin-Echo; TR = 9270 ms; TE = 65 ms, flip angle = 157°, matrix = 256 × 256, FOV 154 × 154 mm^2^, 100 slices and final voxel size = 0.6 × 0.6 × 0.6 mm^3^.

### Imaging analysis

#### Image preprocessing

All preprocessing was implemented using the nipype software package, including FSL, AFNI, and ANTs. To facilitate the extraction of reliable morphometric data all T2-weighted images were skull-stripped and combined (12 degrees-of-freedom [DOF] linear affine transformation) to produce a common reference frame and average T2-weighted template. The T2 template was subsequently non-linearly normalized to the NIMH Macaque Template (NMT v2.0).(Jung et al., 2021; Reveley et al., 2017)

#### Lesion analysis and Region of Interest Definition

The infarcted lesions were segmented using each animal’s 30 day T2-weighted images. All segmentations were performed automatically using FSL’s FAST segmentation and eroded to remove spurious voxels related to edema and cerebrospinal fluid. The resulting segmented lesions were visually inspected by a student and an experienced neurosurgeon (Zhang, Brady, & Smith, 2000). The National Institute of Mental Health Macaque Template (NMT v2.0) normalized Cortical Hierarchy Atlas of the Rhesus Macaque (CHARM) was used to define 278 (139 per hemisphere) cortical regions (Jung et al., 2021; Reveley et al., 2017). For each animal, every region of interest was binarized and the intensity of any voxels that overlapped with the stroke region were subsequently zeroed. This approach effectively eliminates the lesioned voxels for each animal, which could be used later to define the percentage of infarct for each region of interest. Note that we did not use regions from subcortical structures, due to concerns about the anatomically based parcellation of small substructures and decreased signal-to-noise ratio. We also calculated the total stroke volume of each animal after normalization of each animal to the NMT v2.0 template. The total stroke volume of each animal was further decomposed into its constituent grey and white matter components using masks constructed from the NMT v2.0 template. Total stroke, grey and white matter infarct were then subsequently compared between recovery groups to investigate the relationship between lesion size and recovery.

#### Stratifying behavioural recovery

The NHPSS’ aggregate behavioural scores of 11 measures from each animal at 30 days post stroke were submitted to a k-means clustering algorithm to stratify individuals into different recovery groups. The k-means algorithm partitions each animal’s behavioural score into *k* clusters (or recovery groups), such that the within-cluster variance is minimized. The k-means solution was evaluated across values of k ranging from 2 to 6 using the silhouette metric. The k value that returned the highest silhouette score was 2, resulting in a good and poor recovery group.

#### Data Analysis and Predicting Outcomes

The stroke area of each animal was combined to produce a mean and probability map of afflicted cortical brain regions. Each animal was subsequently stratified into their respective good and poor recovery groups. The stroke area and behavioural scores were averaged across animals in each group. We performed non-parametric Mann-Whitney tests to determine significant differences in stroke volume and behavioural scores between recovery groups. We also investigated the relationship between stroke volume and behavioural outcome at 30 days. The relationship between stroke volume and recovery was examined by regressing each animal’s stroke volume with their aggregate NHPSS score.

Our first aim was to determine if the integrity of cortical regions significantly differed between recovery groups. Accordingly, we compared all non-zero voxels (i.e., non-stroke afflicted voxels) within each of the 278 regions of interest between groups using non-parametric Mann-Whitney tests. False Detection Rates (FDR) were applied to the resulting statistics to correct for multiple comparisons. The resulting outcome of these tests is a subset of cortical brain regions that significantly differ in their percent volume remaining as a function of good versus poor recovery groups. There is no way, however, to assign a level of importance to each of these regions in how they relate to good versus poor recovery. Thus, to develop such a measure we attempted to predict behavioural recovery through support vector machine (SVM) binary classification and use the resulting weighting coefficients (one for each region) as an importance map.

To prepare the data for behavioural SVM classification, we restricted our analysis to those regions of interest to the affected hemisphere between the two groups (see above). Non-zero voxel counts for each of the regions of interest were used as predictors within the linear support vector machine. The data was normalized by converting each area’s voxel count into a percentile of the original area’s voxel count. The SVM model used a linear kernel function and a constant cost parameter of 1 to compute the hyperplane that best separated the recovery groups. To verify the generalizability of the data set into two separate recovery groups, we used a leave-one-subject-out cross-validation method, in which the classifier was trained on all animals, except one, which was used as the test case. We performed this procedure iteratively until all animals had been used for classifier testing (i.e., 31 times). The accuracy of classification was determined as the total number of correctly predicted animals divided by the total number of animals. The absolute value of the weighting coefficients for each iteration of the cross-validation procedure were averaged across all iterations to obtain a rank ordered list of the brain regions that contributed most to successful binary classification.

To assess the statistical significance of decoding accuracy with our SVM classifier, we performed permutation testing using non-parametric randomization tests. To do this, for each cross-validation, the labels of the training data (good versus poor recovery) were randomly permuted prior to classifier training, with testing then being performed on the correctly labelled held-out animal’s data. That is, to empirically test the statistical significance of our findings with the true training data, we examined how a model trained on randomized data labels would perform when tested on true labels. We performed this permuted train-and-test procedure until all animals had been held aside for classifier testing (i.e., 31 iterations), and then averaged across iterations to produce a single ‘random’ decoding accuracy. We then performed this entire procedure 1000 separate times, resulting in a null distribution of 1000 mean decoding accuracies. We then found the probability of the true mean decoding accuracy based on its place in a rank ordering of this randomized distribution. Note that the peak percentiles of significance (p < 0.001) are limited by the number of samples producing the randomized probability distribution.

## Data availability

Data is available upon request

## Results

### Trajectory of recovery following stroke

Prior to the MCAO procedure, all the animals (N=31) displayed typical healthy behaviours. The NHPSS evaluation showed a progressive increase in neurological severity score from the first evaluation (24 h post-MCAO), reaching the top score at 4th-day post-MCAO, and then showed a progressive decrease until it became stable at the 28th and 30th-day post-MCAO (Fig 2A).

**Figure 2:**
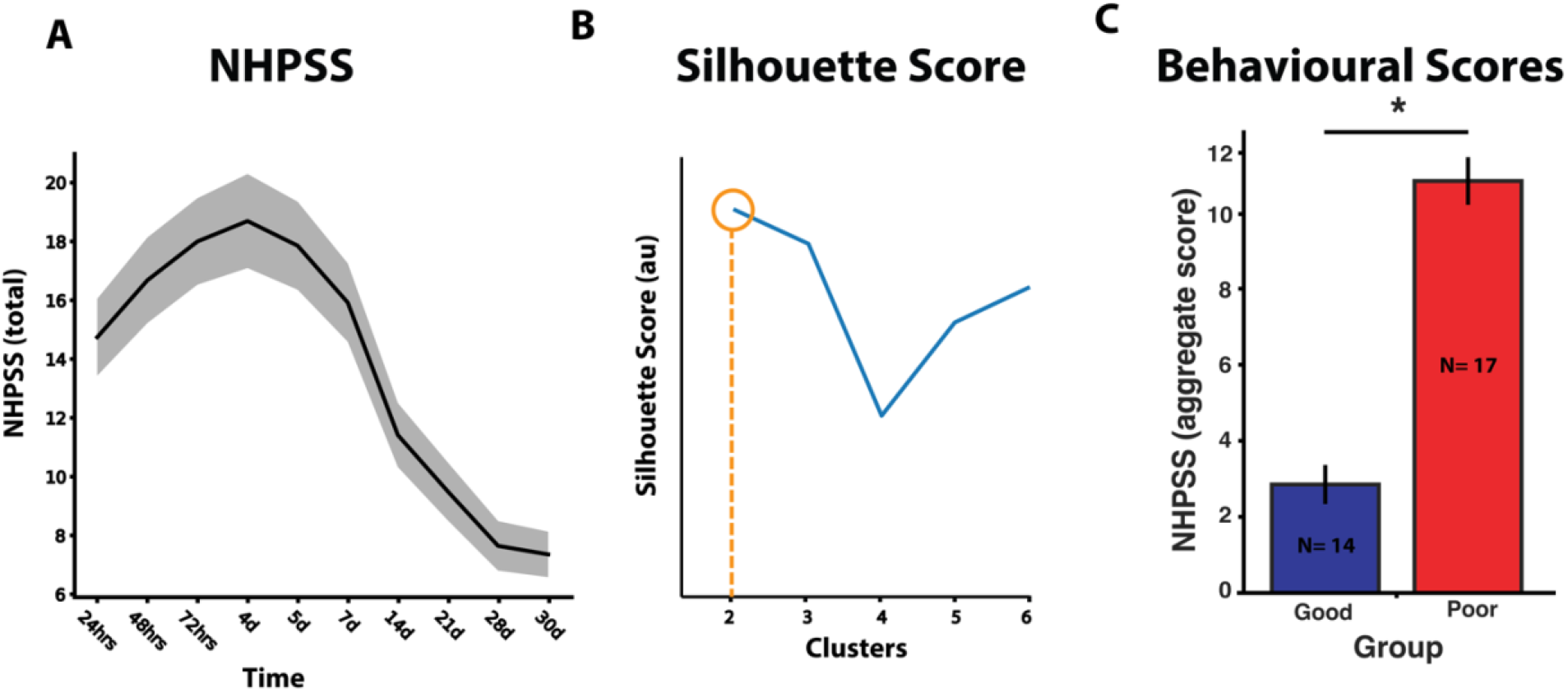
Recovery and behavioural scores. **A)** The average Non-Human Primate Stroke Scale (NHPSS) rating system over 30 days across all animals. **B)** Stratification of animals based on k-means clustering analysis revealed two recovery groups at 30 days post-stroke. The good recovery group consists of 14 animals and the poor recovery group consists of 17 animals. Orange line and circle indicates the optimal stratification of groups. **C)** NHPSS behavioural scores for the good recovery group (blue) and poor recovery group (red). Bars indicate means +/-SEM (* indicates p<0.05)

### Stroke Volume and Behavioural Outcomes

A k-means clustering of NHPSS scores stratified animals into one of two recovery groups, which resulted in a significant difference between the behavioural scores of the two groups (Fig 2C; Mann-Whitney U = 0, p<0.0001). Given that the mean aggregate NHPSS behavioural scores for these groups were 2.96 (SD = 1.71) and 10.58 (SD = 2.11), we interpreted these two groups as indicative of good versus poor recovery, respectively. To investigate the relationship between stroke volume and behavioural recovery, we compared the overall stroke volumes of the two recovery groups. The mean stroke volumes were 6515.07 mm^3^ (SD = 3375.68) and 8146.88 (SD = 3234.93) for the poor and good recovery groups, respectively, and this difference was non-significant (Fig 3A; Mann-Whitney U = 88, p=0.113). Furthermore, the total stroke volume did not predict, across animals, the corresponding NHPSS recovery scores at 30 days (Fig 3B; R^2^ = 0.29), suggesting that overall stroke lesion volume had little bearing on animal recovery. Next, we wondered whether good versus poor group recovery differences would emerge if we considered grey and white matter damage separately. We found that the mean stroke volume was not different between groups within the grey matter 4363.64 (SD = 2176.40) and 5329.65 (SD = 2403.05) for the good and poor recovery groups, respectively. (Fig 3A; Mann-Whitney U = 86, p=0.097). Finally, the remaining non-grey matter stroke volume was also observed to be similar between recovery groups (2151.43 (SD = 1283.76) and 2817.24 (SD = 1194.05); Fig 3A; Mann-Whitney U = 85, p=0.091).

**Figure 3:**
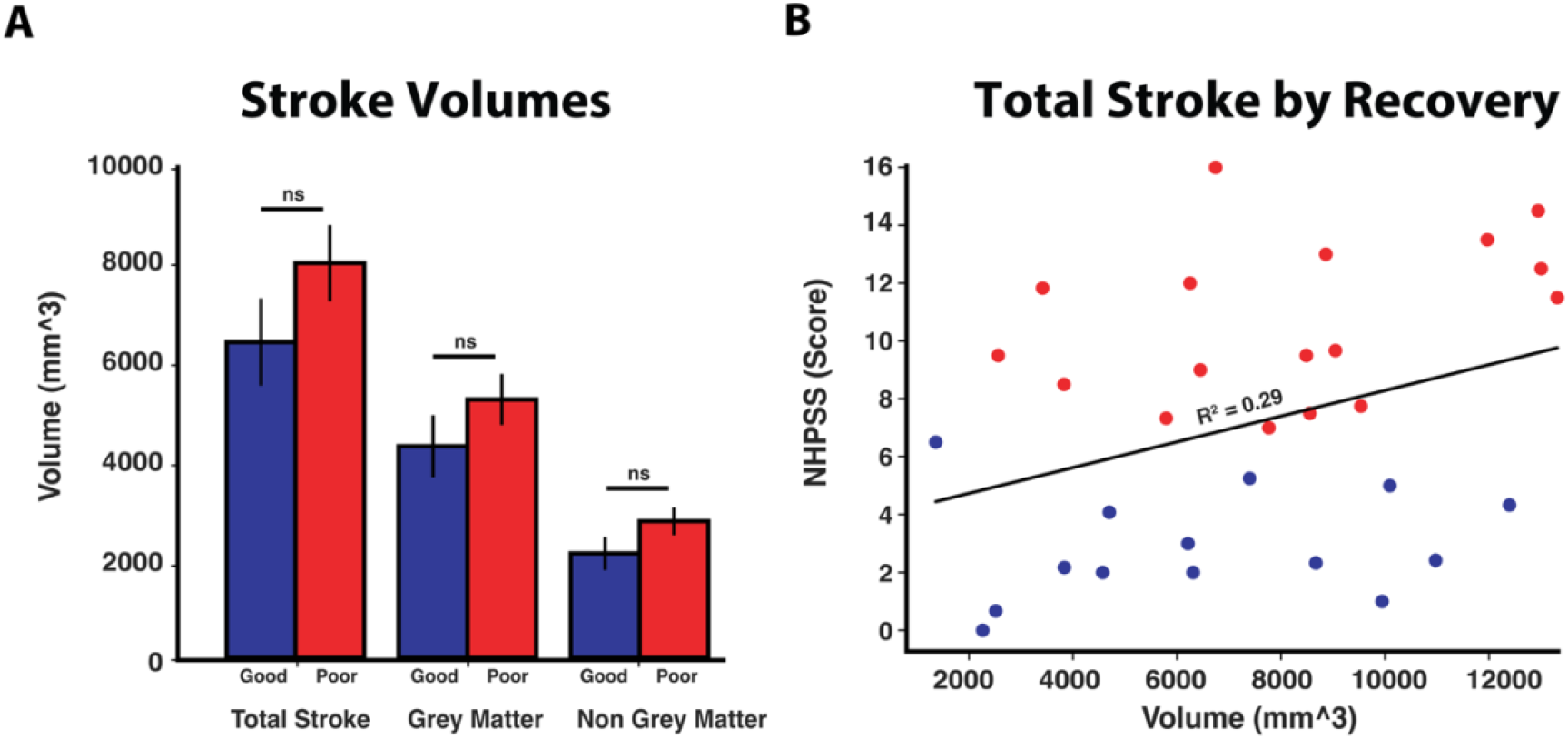
Total stroke volume does not predict recovery outcome. **A)** The total stroke volumes for the good (Blue) and poor (Red) recovery groups. **B)** Scatter plot of all animals’ total stroke volume by their respective recovery scores at 30 days post stroke. Total Stroke volume yielded no correlation with outcomes. Blue denotes the good recovery group and Red denotes the poor recovery group.

### Brain regions differences between groups

Given the lack of differences in stroke volume as a function of recovery group, we next wondered whether these recovery differences were instead attributable to differences in specific brain regions impacted by the stroke. To test this, we examined whether the two recovery groups differed in the percentage of non-stroked cortex for each brain region separately. Across the affected hemisphere, we observed several partially intact brain regions that significantly differed between the good and poor recovery groups (see Fig 4). These regions were mainly within parietal, temporal and frontal cortices. The parietal and temporal regions that displayed differences included the sensory and association regions 3a/b, 7op,7b, TAa (superior temporal gyrus), R (rostral core region of auditory cortex), TPO (Temporal parieto-occipital associated area) and RM (Rostromedial belt region) all of which were salvaged to a greater proportion in the good recovery group compared to the poor recovery group. Our analysis revealed that frontal regions were nearly universally salvaged to a greater extent in the poor recovery group animals compared to the good recovery group animals. Notably, none of the classical motor regions including areas 4,5, and 6 demonstrated significant differences between the recovery groups (Fig 5).

**Figure 4:**
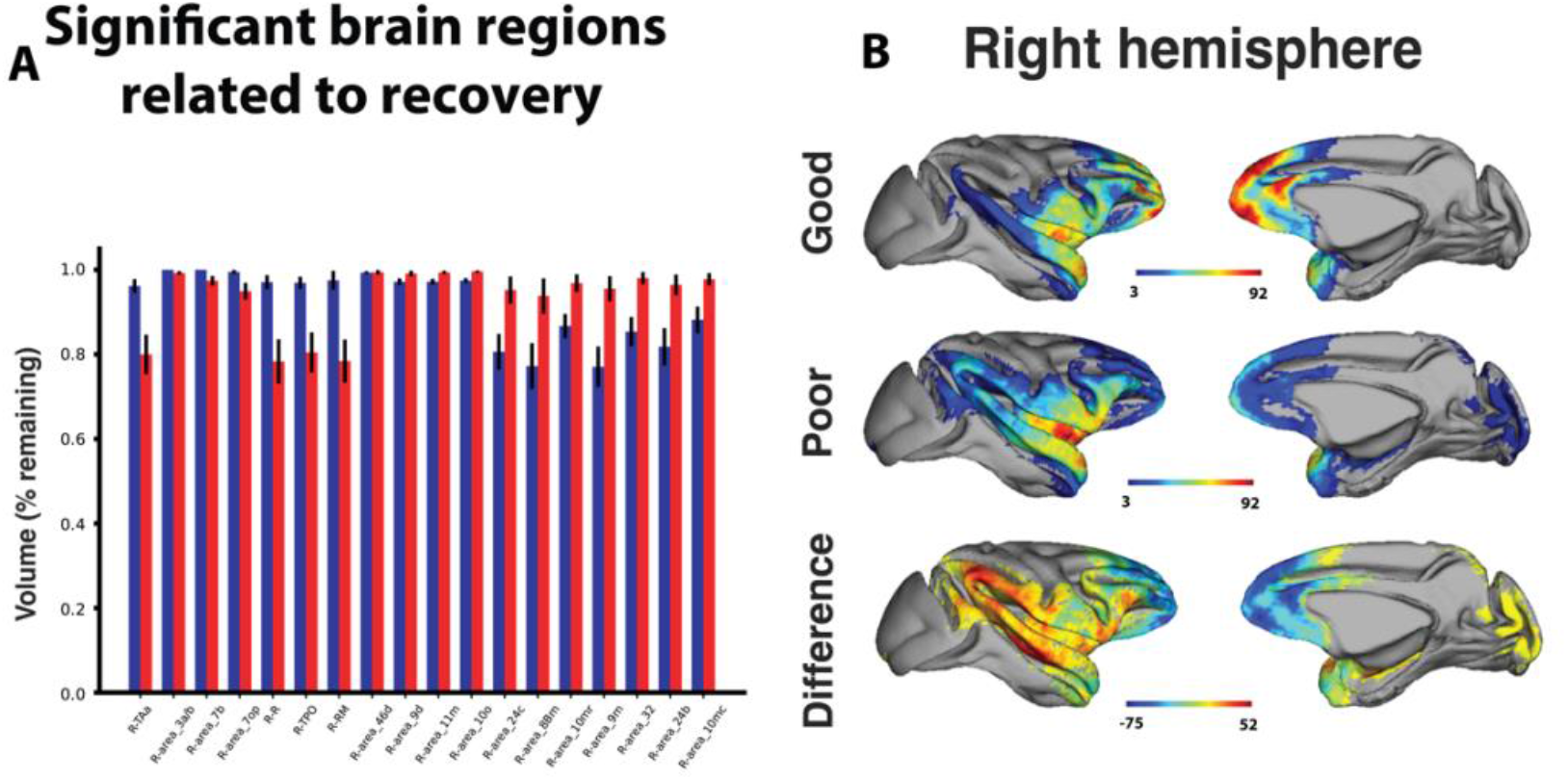
Preservation of areas in parietal temporal cortex is associated with good recovery. **A)** The statistically different areas affected by the strokes in the good recovery group (blue) and poor recovery group (red). The bars denote the mean and SE of the ROI remaining. **B)** Surface projections of the stroke area, represented as a heat map across all animals in the good recovery group (top), poor recovery group (middle), and the difference between the good and poor recovery groups (bottom). Note that for the differences positive values indicate greater areas of salvage for the good recovery group, while negative values indicated greater areas of salvage for the poor recovery group.

**Figure 5:**
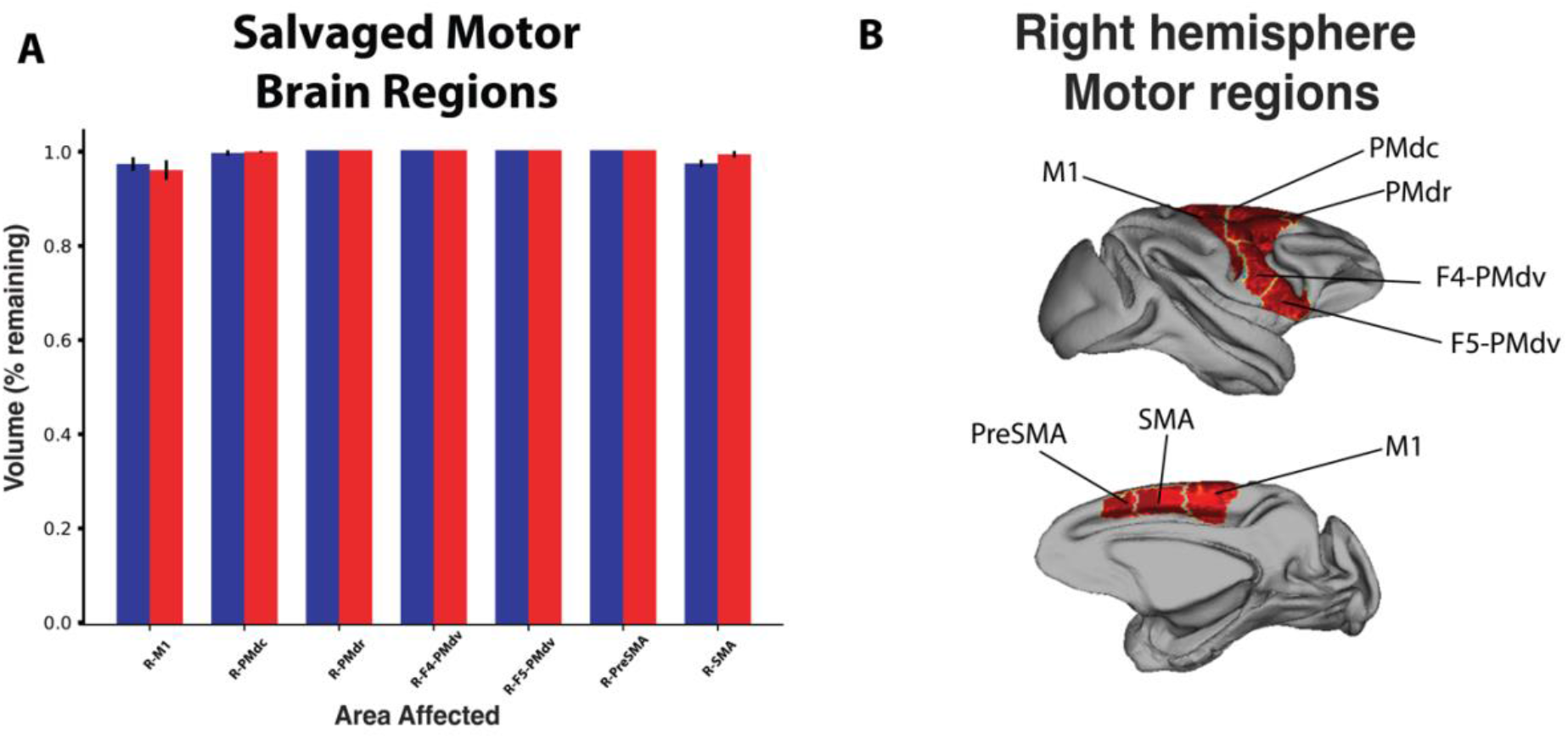
Motor areas are not differentially affected between recovery groups. **A)** Motor areas are largely preserved following the stroke in the good recovery group (blue) and poor recovery group (red). The bars denote the mean and SE of the ROI remaining. **B)** Surface projections of the ROIs presented in panel A.

To independently verify that the brain regions described above are associated with good versus poor recovery, we performed a separate leave-one-subject-out pattern classification analysis wherein, using the proportion of salvaged cortex from all 139 right hemisphere regions (i.e., the stroked hemisphere), we attempted to predict the recovery (good versus poor) of each animal. This analysis revealed significant single animal classification, with a 68% decoding accuracy (see Fig 6). The additional benefit of this approach is that, through the support vector machine classifier, we can use the resulting weighting coefficients (one for each region) from the training data to generate an importance map i.e., delineate the brain regions that have a larger (versus lesser) contribution to accurate decoding. Note that inferring a ranking of importance is not possible with the more conventional univariate approach implemented in Fig. 4.

**Figure 6:**
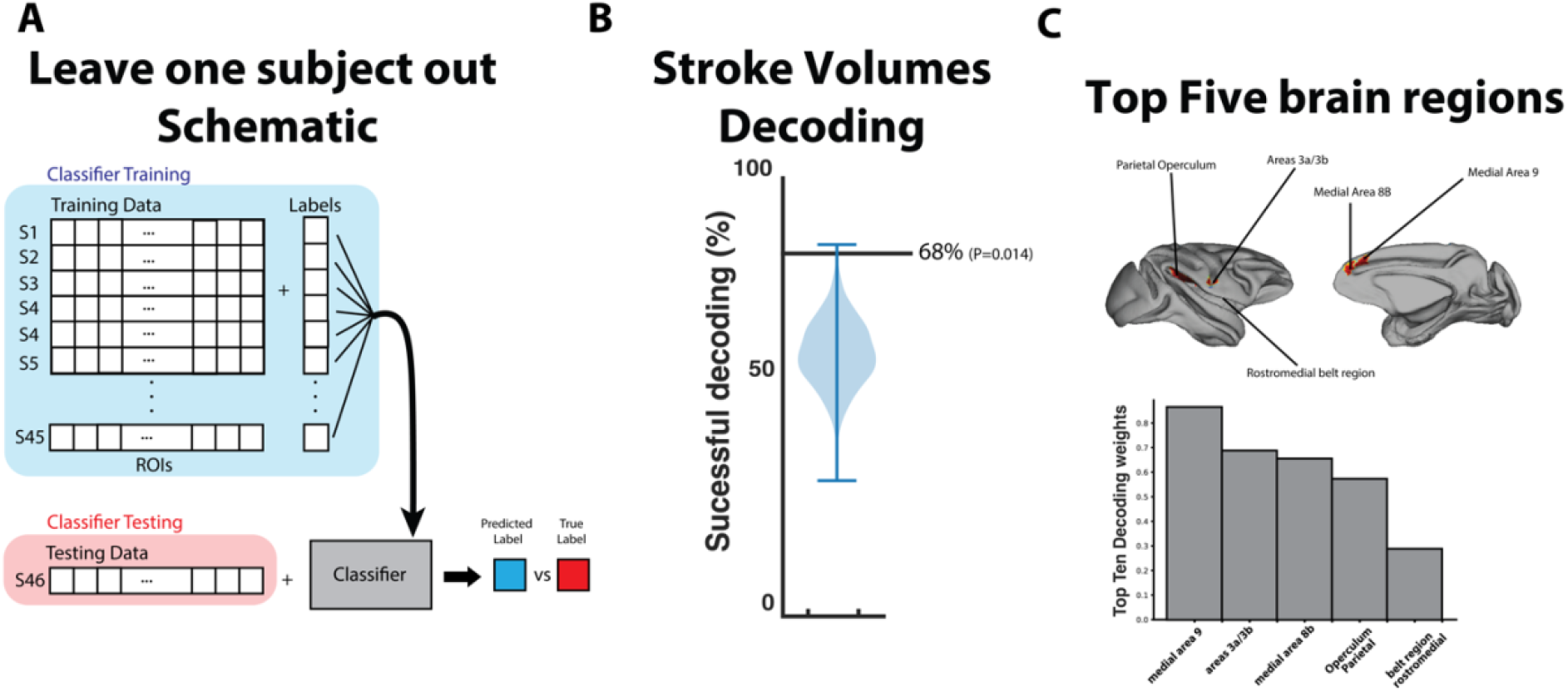
Voxel decoding and top five area contribution. **A)** The decoding schematic for the leave one out machine learning approach taken to determine the decoding accuracy and regional importance to recovery. **B)** the decoding accuracy using the remaining voxels in figure 4A to predict outcomes. **C)** the top five significant areas that determine decoding accuracy displayed on an inflated brain (top) based on their weights from the support vector machine (bottom).

The resulting region-based weights showed that the regions with the largest impact on decoding were medial area 9, areas 3a/b, medial area 8 parietal operculum and rostromedial belt region. Interestingly, medial area 9 and area 3a/b corresponds with significant areas highlighted using the differential lesion analysis between groups above (see Fig 6). The importance of area 8 likely reflect predictors of recovery because these areas tend to be salvaged to a greater extent in the poor group, whereas areas 3a/3b and area 9 tend to be salvaged in the good group.

## Discussion

Here, using structural neuroimaging in 31 macaques, a well-established stroke model, and by exploiting individual differences in animal behavioural spontaneous recovery, we investigated the neural substrates that underlie stroke recovery. We show that differences in outcomes between good and poor recovering animals were tightly linked to the structural integrity of brain areas in sensory and association cortices, specifically those within parietal cortex. Our results also highlight that the most important predictors of spontaneous recovery include key brain regions that have been implicated in multi-modal processing, including somatosensory regions, temporal regions, and posterior parietal regions, which is consistent with recent theories of motor control that posit that successful motor actions are a by-product of intelligent sensory integration (Scott, 2004). Furthermore, motor regions failed to show significant differences between recovery groups. Taken together these findings suggest that the integrity of brain regions involved in integrating sensory information are critical to recovery following stroke.

The course of stroke recovery in humans is highly variable and influenced by several biological and socioeconomic factors. A number of studies have highlighted that major determinants of stroke recovery include previous strokes, co-morbidities, severity of motor deficits, post stroke depression, genetic factors, rehabilitation therapy (eg. timing and dosage), as well as age, gender, race and socioeconomic status (Cramer, 2004, 2008; Hinkle, 2006). The multitude of factors that contribute to heterogeneity in human stroke recovery makes it difficult to disentangle the neural substrates that mediate recovery. The strength of our macaque model is that we have considerably more control over such factors as age, diet, environment, and when assessments and neuroimaging are performed, which creates a well-controlled homogeneous population in which to assess recovery.

Ultimately, the goal of any model is to approximate the human condition as closely as possible. Although primate species can be rather heterogeneous, macaques are phylogenetically close to humans, and closely resemble humans from an anatomical, physiological and behavioural perspective (Cook & Tymianski, 2011, 2012). Macaques possess similar cortical and subcortical functional and structural organization to humans, white:grey matter ratios and vasculature that closely approximates the human cortical anatomy (Cook & Tymianski, 2011, 2012). Importantly, all animals in the current study underwent identical middle cerebral artery occlusions, which closely approximates the most common cerebrovascular impairments in humans (Nogles & Galuska, 2020). The application of the microvascular clip over the middle cerebral artery, proximal to the orbital frontal branch, allows for precise occlusion, which reduces the variability of stroke syndromes typically observed in humans (Cook & Tymianski, 2012). Subsequent removal of the microvascular clip results in the reperfusion of the infarct area, which allows for strict control of the duration of cerebral ischemia. This approach provides a viable, reproducible and realistic non-human primate model of stroke that closely approximates the human brain and pathology.

Beyond the macaque anatomy and physiology, their neurobehaviour closely resembles that of humans (Roitberg et al., 2003). Accordingly, macaques possess the ability to be trained and assessed for cognitive, motor and sensory deficits using standardized tests that are akin to human neurobehavioural tasks, which are the main outcome measure in human clinical trials (Cook & Tymianski, 2012; Roitberg et al., 2003). For example, the NHPSS is a robust and well established neurobehavioural outcome that was developed as an analog to the National Institute of Health Stroke Scale (NIHSS). We employed the NHPSS as a measure of recovery following stroke and stratified animals into groups based on a clustering analysis of the diverse NHPSS measures. Although our unbiased behavioural clustering examined 2-6 clusters, silhouette analysis revealed that optimal partitioning of the data resulted in two recovery groups, good versus poor. The two groups allowed us to directly compare the brain regions that were differentially affected by the MCAO, ultimately allowing us to infer the brain regions underlying good and poor recovery. Importantly, there was no difference in stroke volume measures between animals in the good and poor recovery groups.

Our results illustrated that stroke volume measures, including total stroke volume, grey matter stroke volume and non-grey matter stroke volume, failed to significantly differ between recovery groups or correlate with outcomes. Although some studies have observed correlations between lesion size and functional outcomes, our results are more consistent with previous reports that challenge the notion that lesion size alone is the predominant factor associated with functional recovery (Alexander et al., 2010; Beloosesky, Streifler, Burstin, & Grinblat, 1995; Chen, Tang, Chen, Chung, & Wong, 2000; Miyai, Blau, Reding, & Volpe, 1997; Pantano et al., 1996). Rather, we observed that it was the integrity of key neural substrates that was closely linked to functional recovery. The parietal area represented the bulk of the cortical regions that were significantly preserved to a greater degree in the group with good outcomes compared to those animals with poor outcomes. Specifically, regions 3a/b, 7op, 7b, TAa (superior temporal gyrus), R (rostral core region of auditory cortex), TPO (Temporal parieto-occipital associated area) and RM (Rostromedial belt region) were all found to be salvaged to a greater degree in animals in the good group compared to those animals in the poor group. These areas represent secondary sensory and association regions that all play an intimate role in sensory processing and have rich projections to various sensory and motor regions throughout the brain (Carmichael & Price, 1996; Cipolloni & Pandya, 1999; Felleman & Van Essen, 1991; Godschalk, Lemon, Kuypers, & Ronday, 1984). For example, neurons within area 3a/b are well known to play a critical role in somatosensation and sensory integration. This area has been shown to produce complex responses to a variety of sensory stimuli and have rich projections to orbitofrontal and motor regions (Carmichael & Price, 1996; Cipolloni & Pandya, 1999; Felleman & Van Essen, 1991; Godschalk et al., 1984). Posterior parietal regions, including area 7, dorsal prelunate and LIP are fundamental structures in visuo-motor integration and control (Colby, Duhamel, & Goldberg, 1996; Culham & Valyear, 2006; Iacoboni, 2006). For example, disruption of posterior parietal cortex results in apraxia or altered reaching movements that result in the inability to perform movement corrections (Della-Maggiore, Malfait, Ostry, & Paus, 2004; Desmurget et al., 1999; Sathian et al., 2011). Similarly, temporal regions represent areas that have been heavily implicated as hubs for auditory processing and play an important role in spatial localizations and possess extensive cortical and subcortical projections (Plakke & Romanski, 2014; Roumazeilles et al., 2020). Recent evidence suggests that neural activity within these regions are modulated during motor planning, suggesting that disruption of areas within temporal regions could substantially impair motor planning and execution (Gale et al., 2021).

Our classification results highlighted the importance of sensory areas and association cortices as key components in recovery following stroke. Although parietal areas were largely responsible for the differences between recovery groups, our decoding analysis also suggested that the integrity of medial area 9 and areas 3a/3b appeared to play a key role in the good recovery group compared to the poor recovery group. These areas play key roles in a variety of functions such as spatial memory, recognition, and sensory integration (Borich, Brodie, Gray, Ionta, & Boyd, 2015; Kaas, Nelson, Sur, Lin, & Merzenich, 1979). For example, area 3 has rich connections to a variety sensory and motor regions throughout the brain. Damage to area 3 has been shown to produce tactile deficits, including numbness and tingling which appear to impair successful motor control. Overall, the results of our study suggest that sensory processing, and the brain regions that mediate them play a critical role in recovery. The integrity of these sensory areas likely contributes to the successful coordination of motor actions, which are ultimately responsible for improved functional outcomes.

### Methodological Considerations

There are a few methodological considerations to account for in the present study. It is important to acknowledge some of the challenges associated with interpreting the findings from lesions studies. First and foremost, many lesion studies assume consistent brain modularity or discrete anatomical units that are tightly linked to specific functions (Rorden & Karnath, 2004). However, increasing evidence suggests that many neural functions are carried out in parallel or in a distributed manner (Farah, 1994). For example, action control is thought to be supported through frontoparietal networks, with specialized pathways dedicated to motor-related functions like reaching, grasping and eye movements (Haar & Donchin, 2020; Shadmehr & Krakauer, 2008). Thus, it may not be obvious how damage to one region within a distributed circuit affects the entire network.

Next, the approach used here makes a large assumption that superimposition of lesions will always identify the functional brain regions and neural substrates affected. However, there is considerable variability in the brains of individuals and different regions can show different functional responses to damage. Our analysis simply highlights regions implicated in recovery and makes no comment on potential remapping of affected brain regions, which has been well documented in literature (Murphy & Corbett, 2009).

It is important to note that we did not collect any diffusion tensor imaging and thus cannot comment on the individual tracts affected by the strokes in the present study. However, it should be noted that the data suggests that it does not appear that recovery was at all dependent on white matter damage since non-grey matter stroke was similar between recovery groups. Finally, the present study was limited to assessments of macroscopic brain structure, and not function. Thus, we can’t comment on how functional coupling between regions may have preserved or impacted the patterns of damage observed. Correspondingly, we did not quantify or characterize any macroscopic rewiring or molecular pathways. However, these are clear future directions of study.

## Conclusions

Here we show that differences in outcomes between good and poor recovering animals were tightly linked to the structural integrity of brain areas in sensory and association cortices. Furthermore, we highlight that the integrity of parietal regions plays a key role in predicting outcomes following stroke. Taken together, our results suggest that functional outcomes may be mediated by the preservation of sensory regions as they may underlie successful motor control.

## Abbreviations

ACA: Anterior Cerebral Artery
ICA: Internal Carotid Artery
MCA: Middle Cerebral Artery
PCA: Posterior Cerebral Artery
MCAO: Middle Cerebral Artery Occlusion
NHPSS: Non-Human Primate Stroke Scale

